# Neuromodulation of zebrafish primary motoneuron firing is shaped by developmental changes in the M-current

**DOI:** 10.1101/2025.10.06.680750

**Authors:** Stephanie F. Gaudreau, Tuan V. Bui

## Abstract

Movements during development are refined through the ongoing maturation of the spinal circuits that mediate them. In many vertebrates, including zebrafish, this maturation process involves neuromodulators such as acetylcholine, serotonin, and dopamine; however, the targets of this neuromodulation remain largely unknown. Recent work has revealed distinct developmental dynamics of the non-inactivating subthreshold potassium current – the M-current – in primary motoneurons of larval zebrafish. Considering that neuromodulators play a role in the maturation of locomotor control in larval zebrafish, we asked whether neuromodulators might target the M-current in primary motoneurons during development. Our patch-clamp experiments in primary motoneurons of zebrafish aged 3 to 5 days post-fertilization (dpf) reveal distinct modulation of the M-current by serotonin and acetylcholine that is age-dependent. Our data demonstrates an inhibitory influence of serotonin signaling via 5HT_1A_ receptors that promotes repetitive firing in primary motoneurons specifically at 3 dpf. We also show that acetylcholine, likely via M2/M4 receptors, enhances the M-current and limits repetitive firing in primary motoneurons but does so only after 3 dpf. Modulation of the M-current in primary motoneurons was not observed across all neuromodulators as dopamine had no effect at any age. Considering that the M-current transiently peaks at 3 dpf and is reduced at 4 and 5 dpf, our findings suggest that the developmental changes in this current can shape how neuromodulators modulate firing properties of primary motoneurons.

**Key points:**

- The amplitude of the M-current is subject to modulation by several neuromodulators such as acetylcholine and serotonin
- Serotonin via 5HT_1A_ inhibits the M-current in primary motoneurons at 3 dpf
- Muscarinic agonist enhances the M-current in primary motoneurons after 3 dpf
- Neuromodulation of motoneuron firing properties by 5-HT_1A_ and muscarinic agonist is consistent with modulation of the M-current

## INTRODUCTION

Movement is generated by specialized neuronal networks located within the spinal cord. These spinal motor circuits must undergo progressive maturation during development to allow for increasingly refined and complex movements to be made. The exact changes spinal motor circuits undergo during maturation are not fully known but there is evidence supporting a role for neuromodulation in motor maturation. For example, both dopamine (DA) and serotonin (5-HT) have been shown to elicit air-stepping in spinally-transected neonatal rats, supporting their contributions to promoting hindlimb stepping (McEwen *et al*., 1997). In the developing frog *Xenopus laevis,* 5-HT and noradrenaline have contrasting and coordinated influences on separate spinal locomotor circuits (Rauscent *et al*., 2009). While effects to motor output by neuromodulatory systems during development have been described, little is known about the targets of this neuromodulation and the mechanisms by which they shape motor maturation.

The zebrafish is particularly well-suited for the study of motor maturation as their rapid development comprises stereotyped transitions in the movements they produce. Changes in motor behaviour can be readily linked to changes in underlying spinal circuitry. Indeed, DA and 5-HT have both been shown to have important influences on locomotor output during zebrafish development. DA signaling in the spinal cord mediates the important locomotor transition from 3 days post-fertilization (3 dpf) burst swimming to 4 dpf beat and glide swimming (Lambert *et al*., 2012). It has also been shown to control the excitation of motoneurons (Jha & Thirumalai, 2020).

5-HT has been shown to limit the duration of periods of inactivity between successive swimming episodes in developing zebrafish (Brustein *et al*., 2003). While these findings have proven important to our understanding of how supraspinal regions mediate flexible control over zebrafish spinal locomotor circuits in the context of development, the mechanisms by which these neuromodulators exert their influence on spinal neurons are unknown.

We have recently identified the presence of the non-inactivating subthreshold potassium current known as the M-current (*I*_M_) in primary motoneurons (pMNs) of larval zebrafish. We show that *I*_M_’s influence on the control of excitability and regulation of repetitive firing changes during early zebrafish development, peaking at 3 dpf (Gaudreau & Bui, 2024). This coincides with the time when larvae are largely inactive and exhibit burst swimming, an immature form of swimming. By 4 dpf, their mature locomotor pattern – known as beat and glide swimming – emerges, and they swim much more frequently (Buss & Drapeau, 2001; Drapeau *et al*., 2002).

Could *I*_M_ be a target for neuromodulation during development to facilitate motor maturation? As zebrafish transition from their embryonic stage characterized primarily of large amplitude and reflexive pMN-driven movements to their larval stage encompassing predominantly low amplitude secondary motoneuron (sMN) driven swimming, we posit that *I*_M_ in pMNs may be an important target for neuromodulation as pMN recruitment is restricted to specific movements (Liu & Westerfield, 1988; Menelaou & McLean, 2012; Wen *et al*., 2020).

In this study, we use whole-cell electrophysiology to investigate how neuromodulators influence the properties of *I*_M_ in pMNs of larval zebrafish from 3 dpf to 5 dpf. Interestingly, we find that muscarine, a known inhibitor of *I*_M_ in other systems (Brown & Adams, 1980), does not inhibit *I*_M_ in pMNs of larval zebrafish and instead acts as an *I*_M_ enhancer. We find that enhancement of *I*_M_ by muscarine is age-dependent and coincides with ages where the current is relatively smaller. On the other hand, we observed inhibition of *I*_M_ by 5-HT via 5-HT_1A_ receptors. This inhibition is also age-dependent and occurs when *I*_M_ is at its largest within the developmental timeframe studied, namely at 3 dpf. Furthermore, we demonstrate that muscarine and a 5-HT_1A_ agonist modulate pMN firing in a manner consistent with their respective modulations of *I*_M_.

## METHODS

### Animal Care

All experiments were performed in accordance with the protocol approved by the University of Ottawa’s Animal Care Committee (BL-4416). As described previously (Gaudreau & Bui, 2025), adult zebrafish are maintained at 28.5°C with a 14 hour on/10 hour off light cycle, with lights off at 11PM and on at 9AM. Embryos are fertilized between 9AM-10AM and stored in embryo medium at 28.5°C until use for experimentation at 3 to 5 dpf.

### Preparation for electrophysiology

Larval zebrafish were anesthetized in 0.02% tricaine (MS-222, Aqualife TMS; Syndel Laboratories) before being pinned down through the notochord onto a Sylgard (Dow Corning) coated dish using pins made of Tungsten wire (0.25 mm). Fish were spinalized at the level of segments 2-3 using fine surgical scissors, and the skin is then peeled back between the two pins using fine forceps. Next, muscles were dissociated by bathing larvae in 1 mg/mL collagenase (Millipore Sigma; incubation times: 10 minutes at 3 dpf, 12 minutes at 4 dpf, and 15 minutes at 5 dpf) to facilitate muscle removal. The collagenase solution is rinsed out 4-5 times with artificial cerebrospinal fluid (aCSF) containing: 134 mM NaCl, 2.9 mM KCl, 1.2 mM MgCl_2_, 2.1 mM CaCl_2_, 10 mM dextrose, and 10 mM HEPES (pH of 7.8 adjusted with NaOH and osmolarity between 280-290 adjusted with sucrose). Removal of muscles was done using suction through a wide-bored glass capillary whose tips were gently broken such that the tip was the appropriate size for muscle removal. To enable whole-cell patch-clamp electrophysiology, muscles overlaying the targeted spinal segments were removed.

### Whole-cell patch-clamp electrophysiology

d-tubocurarine was added to the aCSF (10 μM; Millipore Sigma, catalog # 93750) to prevent muscle contractions. Electrodes for whole-cell patch-clamp experiments are made using borosilicate glass capillaries (Sutter Instrument; BF150-110-10) with typical tip resistances ranging from 5 to 7 MΩ. Intracellular recording solution used contained the following: 16 mM KCl, 116 mM K-gluconate, 4 mM MgCl_2_, 10 mM HEPES, 10 mM EGTA, and 4 mM Na_2_-ATP, adjusted to a pH of 7.2-7.3 with KOH. Osmolarity was adjusted to 270-280 mOsm with water. The initial descent of the electrode towards the spinal cord involved applying slight positive pressure.

Upon proximity to a pMN (targeted based on their distinctively large soma and mid-dorsoventral positioning), positive pressure was released to form a gigaohm (GΩ) seal. Pulses of negative pressure were used to break through the membrane. Membrane holding potential was quickly set to –65 mV and compensation of both fast and slow capacitance components was applied. Series resistance was routinely compensated for at 80% or higher. Values reported are not liquid junction potential corrected. Sulforhodamine B was added to the intracellular recording solution (0.1%; Millipore Sigma) to identify pMNs by visualization of their axon projection patterns. Electrical signals were amplified and filtered at 10 kHz or 20 kHz with a Multiclamp 700B (Axon Instruments, Molecular Devices),and digitized with a Digidata 1550 (Molecular Devices). A HumBug Noise Eliminator (Quest Scientific) attenuated 50/60 Hz electrical noise.

### Voltage-clamp protocols and current estimations

*I*_M_ was revealed through the standard *I*_M_ deactivation protocol (Verneuil *et al*., 2020; Sharples *et al*., 2023) in voltage-clamp mode. The protocol consists of holding the neuron at −10 mV and introducing a series of hyperpolarizing voltage steps lasting 1 second each to deactivate *I*_M_. The resulting loss of outward current caused by *I*_M_ deactivation can be used to estimate the amplitude of *I*_M_. This calculation was made by taking the difference between the peak of the initial current response and the current at steady-state at the end of the step.

### Current-clamp protocols and firing frequency measurements

500-ms current steps were applied to neurons initially held at −65 mV to calculate firing frequency. Rheobase was defined as the minimum current amplitude required to elicit an action potential. During a 500-ms current injection equal to 2x rheobase, the average instantaneous frequency of the last 5 pairs of action potentials was used to estimate steady-state firing frequency.

### Pharmacology

Dopamine hydrochloride (3548, Tocris), serotonin hydrochloride (3547, Tocris), muscarine iodide (3074, Tocris), 5-HT_1A_ agonist 8-OH-DPAT (1080, Tocris), M2/M4 agonist oxotremorine-M (O100, Millipore Sigma), and 5-HT_2A_ agonist TCB-2 (2592, Tocris) were dissolved in water before final dissolution in aCSF of the recording bath. All drugs were introduced to the recording bath 5-10 minutes prior to recordings.

### Data analysis and statistical analysis

We used the open-source pyABF python package in Spyder (version 5.1.5) to import and read electrophysiological data saved as .abf files. Python (version 3.9.12) scripts tailored to each type of recording enabled supervised, automated analysis of recordings. Prism by GraphPad (Version 10.5.0 (673) was used for statistical analyses and generation of graphs. The Shapiro-Wilk test was used to test all datasets for normality. For all tests, significance stars are displayed on graphs.

## RESULTS

### Muscarine enhances I_M_ in pMNs at 4 and 5 dpf

The discovery of *I*_M_ and its appellation originated from its inhibition by muscarine (Brown & Adams, 1980). Since our previous work identified *I*_M_ in pMNs of larval zebrafish (Gaudreau & Bui, 2024), we sought to investigate the potential for muscarinic inhibition of *I*_M_ in these neurons (**Fig. 1**). As described previously (Verneuil *et al*., 2020; Sharples *et al*., 2023; Gaudreau & Bui, 2024), we utilized the standard *I*_M_ voltage-clamp deactivation protocol to reveal its electrophysiological signature in current responses. Comparison of effects were made between three entirely distinct sample groups of pMNs: control pMNs and two groups of pMNs having been exposed to either 10 µM muscarine or 100 µM muscarine for several minutes prior to all recordings. Because of differences observed in the magnitude of *I*_M_ in pMNs during early zebrafish development, we compared the effects of muscarine at 3, 4, and 5 dpf. It is important to note that all data obtained from the control groups of pMNs presented here has been presented in prior work (Gaudreau & Bui, 2024). At 3 dpf, exposure to 10 µM muscarine and 100 µM muscarine did not alter the current-voltage relationship of *I*_M_ relative to control (**Fig. 1*A-B***). No significant differences in peak amplitude of *I*_M_ (control: 42.88 ± 18.44 pA; 10 µM muscarine: 50.56 ± 21.87 pA; 100 µM muscarine: 29.96 ± 18.99 pA; **Fig. 1*C***), *I*_M_ activation voltage (control: −54.55 ± 5.68 mV; 10 µM muscarine: −58.00 ± 9.19 mV; 100 µM muscarine: −49.00 ± 10.22 mV; **Fig. 1*D***), or the voltage at which *I*_M_ reaches half its maximal amplitude (*V*_1/2_) (control: −47.64 ± 4.31 mV; 10 µM muscarine: −51.52 ± 6.99 mV; 100 µM muscarine: −43.98 ± 8.95 mV; **Fig. 1*E***) were observed between either 10 µM or 100 µM muscarine and controls. At 4 dpf, while 100 µM muscarine did not alter any of the properties of *I*_M_, 10 µM muscarine enhanced *I*_M_ (**Fig. 1*F-J***). The average amplitude of *I*_M_ across voltages was significantly increased in pMNs exposed to 10 µM muscarine compared to controls (**Fig. 1*F-G***). The peak amplitude of *I*_M_ was also significantly increased in pMNs exposed to 10 µM muscarine compared to control pMNs (control: 24.08 ± 10.91 pA; 10 µM muscarine: 42.86 ± 22.62 pA; 100 µM muscarine: 26.95 ± 15.81 pA; **Fig. 1*H***). The activation voltage of *I*_M_ (control: −45.63 ± 10.63 mV; 10 µM muscarine: −56.50 ± 3.38 mV; 100 µM muscarine: −49.09 ± 12.61 mV; **Fig. 1*I***) and *V*_1/2_ (control: −41.90 ± 6.74 mV; 10 µM muscarine: −50.71 ± 4.30 mV; 100 µM muscarine: −42.06 ± 12.81 mV; **Fig. 1*J***) in pMNs were significantly hyperpolarized by 10 µM muscarine but remained unaltered by 100 µM muscarine compared to controls at 4 dpf. By 5 dpf, the average amplitude of *I*_M_ across voltages remains increased in pMNs exposed to 10 µM muscarine relative to control pMNs (**Fig. 1*K-L***) yet the peak amplitude is not significantly different between these groups (control: 28.63 ±11.56 pA; 10 µM muscarine: 39.59 ± 14.75 pA; **Fig. 1*M***). The current-voltage relationship of pMNs exposed to 100 µM muscarine compared to controls remained unchanged (**Fig. 1*K***) as did the peak amplitude of *I*_M_ (100 µM muscarine: 24.90 ± 10.53 pA; **Fig. 1*M***). The activation voltage of *I*_M_ was hyperpolarized by 10 µM but not 100 µM muscarine (control: −49.09 ± 7.06 mV; 10 µM muscarine: −56.36 ± 3.93 mV; 100 µM muscarine: −50.45 ± 6.11 mV; **Fig. 1*N***). Finally, the *V*_1/2_ of *I*_M_ in pMNs exposed to either 10 or 100 µM muscarine was unchanged relative to controls (control: −47.79 ± 3.37 mV; 10 µM muscarine: −49.91 ± 2.94 mV; 100 µM muscarine: −41.43 ± 9.71 mV; **Fig. 1*O***). Overall, these data suggest that muscarine enhances *I*_M_ at a concentration of 10 µM, but not 100 µM, at 4 dpf and 5 dpf, but not 3 dpf.

**Figure 1.**
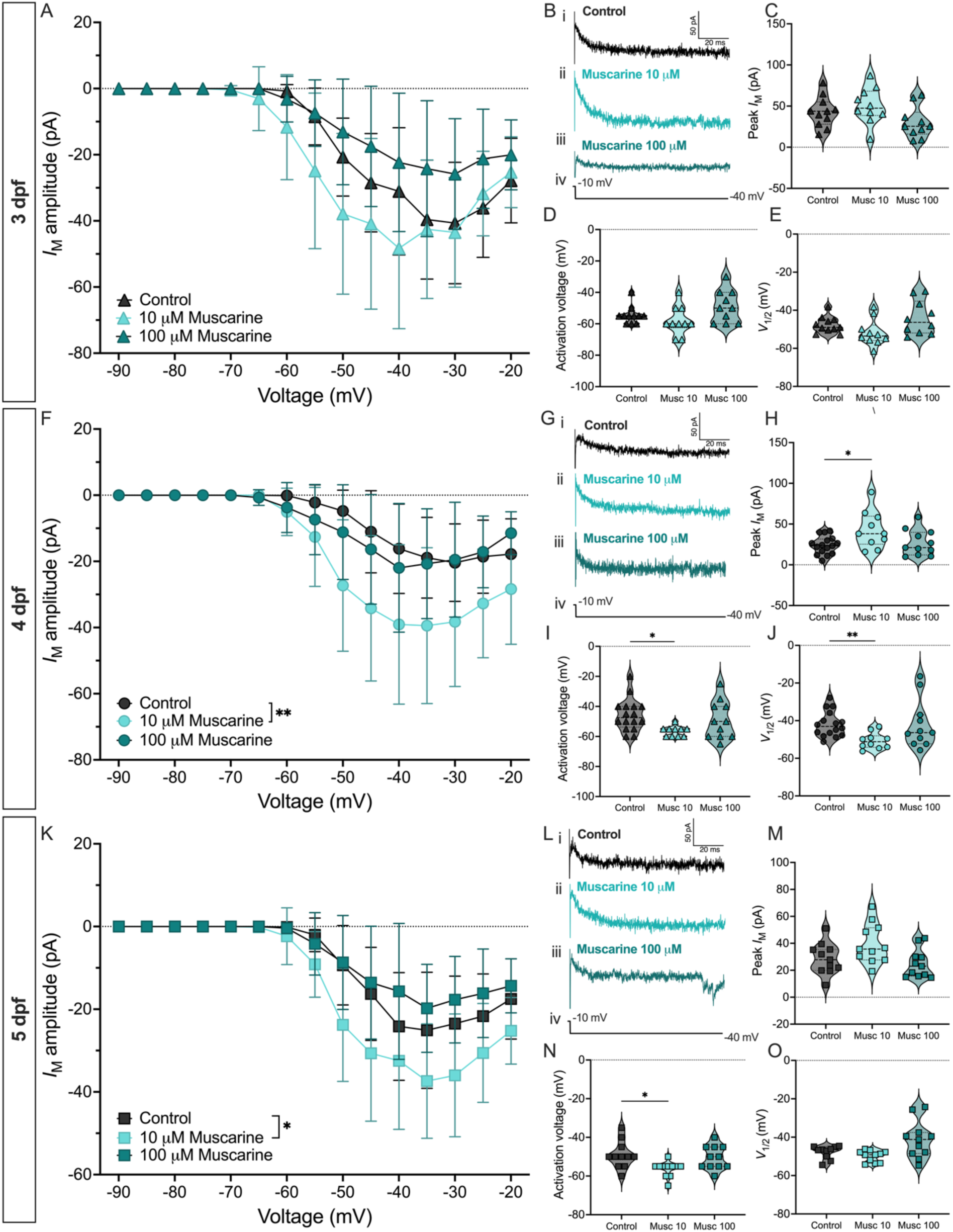
Muscarine enhances *I*_M_ in pMNs of developing zebrafish after 3 dpf. ***A***, Current-voltage relationship of *I*_M_ in three groups of 3 dpf pMNs: control (*n* = 11, *N =* 6), 10 µM muscarine (Musc 10; *n* = 10, *N* = 4), 100 µM muscarine (Musc 100; *n* = 10, *N* = 4). ***B***, Examples of current responses from (*i*) control, (*ii*) 10 µM muscarine, and (*iii*) 100 µM muscarine pMNs in response to (*iv*) a −30 mV hyperpolarizing voltage step at 3 dpf. ***C***, Peak amplitude of *I*_M_ across treatment groups at 3 dpf. ***D***, Activation voltage of *I*_M_ across treatment groups at 3 dpf. ***E***, Voltage at which *I*_M_ reaches half its maximal amplitude (*V*_1/2_) across treatment group at 3 dpf. ***F***, Current-voltage relationship of *I*_M_ in three groups of 4 dpf pMNs: control (*n* = 16, *N =* 6), 10 µM muscarine (*n* = 10, *N* = 3), 100 µM muscarine (*n* = 11, *N* = 5). ***G***, Examples of current responses from (*i*) control, (*ii*) 10 µM muscarine, and (*iii*) 100 µM muscarine pMNs in response to (*iv*) a −30 mV hyperpolarizing voltage step at 4 dpf. ***H***, Peak amplitude of *I*_M_ across the treatment groups at 4 dpf. ***I***, Activation voltage of *I*_M_ across treatment groups at 4 dpf. ***J***, Voltage at which *I*_M_ reaches half its maximal amplitude (*V*_1/2_) across treatment group at 4 dpf. ***K***, Current-voltage relationship of *I*_M_ in three groups of 5 dpf pMNs: control (*n* = 11, *N =* 5), 10 µM muscarine (*n* = 11, *N* = 3), 100 µM muscarine (*n* = 11, *N* = 4). ***L***, Examples of current responses from (*i*) control, (*ii*) 10 µM muscarine, and (*iii*) 10 µM muscarine pMNs in response to (*iv*) a −30 mV hyperpolarizing voltage step at 5 dpf. ***M***, Peak amplitude of *I*_M_ across the treatment groups at 5 dpf. ***N***, Activation voltage of *I*_M_ across treatment groups at 5 dpf. ***O***, Voltage at which *I*_M_ reaches half its maximal amplitude (*V*_1/2_) across treatment group at 5 dpf (control: *n* = 10, *N* = 5; rest of sample sizes same as in ***K***). *Statistical analysis,* (***A***, ***F***, ***K***) mixed-effects analysis, (***D***, ***I-J***, ***N-O***) Kruskal-Wallis test with Dunn’s test for multiple comparisons, and (***C***, ***E***, ***H****, **M***) ordinary one-way ANOVA with Dunnett’s test for multiple comparisons: * *P* < 0.05, ** *P* < 0.01; otherwise, not statistically significant.

### Acetylcholine signaling via M2 & M4 receptors enhances I_M_ in pMNs at 4 dpf

Having revealed that muscarine enhances *I*_M_ in pMNs at 4 and 5 dpf, we wanted to identify the receptors via which acetylcholine (ACh) is exerting its influence. We bath applied 20 µM oxotremorine-M (Oxo-M), a broad muscarinic agonist with some reports of higher affinity for M2 and M4 receptors (Bevan, 1984; Caulfield & Brown, 1991), to pMNs and observed how the properties of *I*_M_ changed relative to control pMNs at 4 dpf. Oxo-M enhanced the average amplitude of *I*_M_ across voltages (**Fig. 2*A-B***) as well as the peak amplitude of *I*_M_ (control: 24.08 ± 10.91 pA; Oxo-M: 65.84 ± 18.92 pA; **Fig. 2*C***). Neither the activation voltage (control: −45.63 ± 10.63 mV; Oxo-M: −51.67 ± 4.33 mV; **Fig. 2*D***) nor *V*_1/2_ (control: −41.90 ± 6.75 mV; Oxo-M: −46.43 ± 3.91 mV; **Fig. 2*E***) of *I*_M_ was altered in pMNs at 4 dpf. Overall, these results demonstrate that ACh signaling via M2/M4 receptors enhances *I*_M_ in pMNs at 4 dpf.

**Figure 2.**
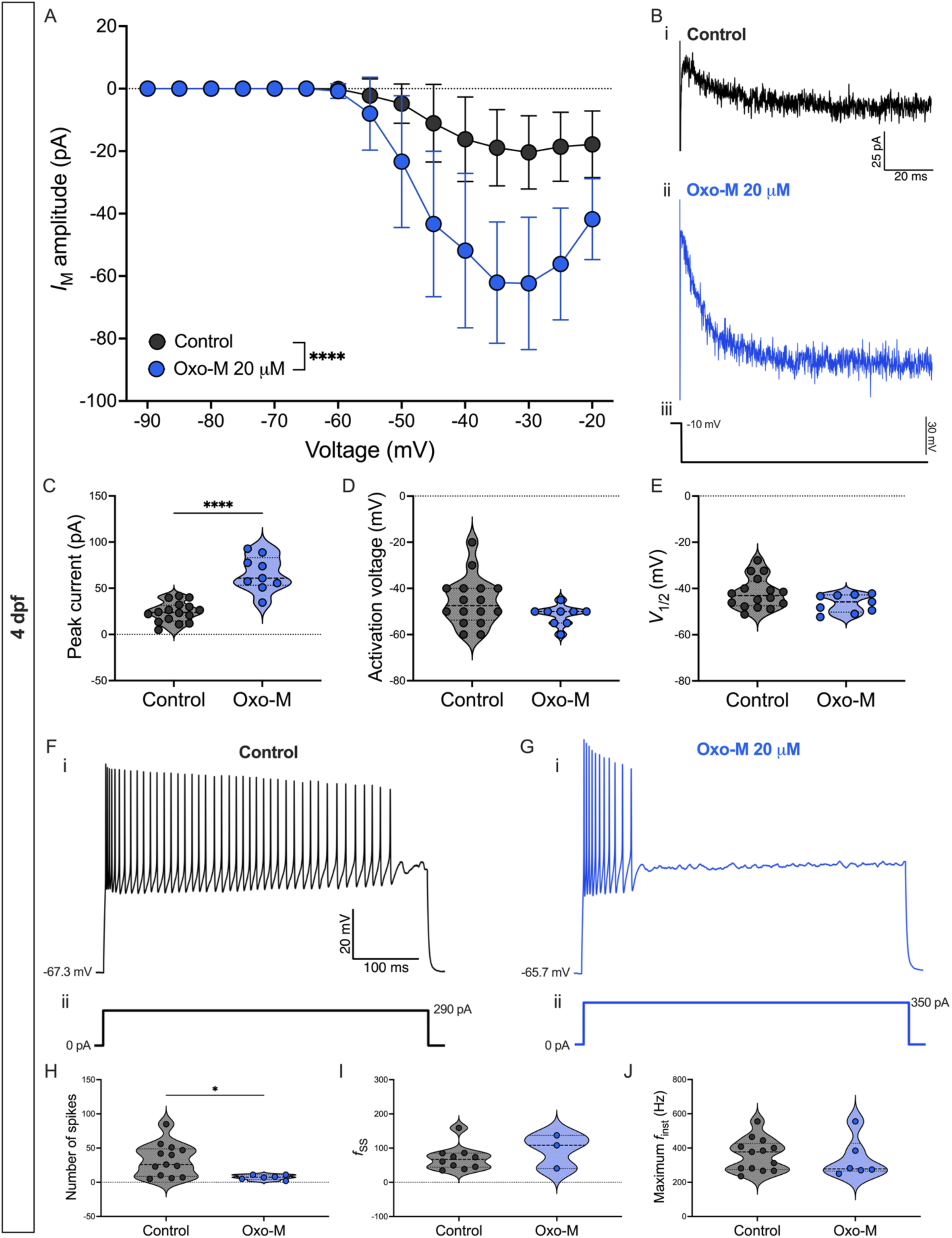
mAChR agonist enhances *I*_M_ in pMNs at 4 dpf. ***A***, Current-voltage relationship of *I*_M_ in control pMNs (black) and those exposed to 20 μM oxotremorine-M (Oxo-M) at 3 dpf. ***B***, Example current responses of (***i***) a control pMN (*n* = 16, *N* = 7) and (***ii***) a pMN exposed to 20 μM Oxo-M (*n* = 9; *N* = 4) in response to (***iii***) a negative 30 mV step from −10 mV holding during standard *I*_M_ deactivation protocol. ***C***, Peak *I*_M_ amplitude. ***D***, Activation voltage of *I*_M_. ***E***, Voltage at which *I*_M_ reaches half its maximal amplitude (*V*_1/2_). ***F***-***G***, Representative current-clamp responses of (***Fi***) control pMNs and ***Gi***) those exposed to 20 μM Oxo-M in response to (***ii***) a current injection equal to 2x rheobase. ***H***, Number of spikes generated in response to a 2x rheobase current injection (control: *n* = 13, *N* = 5; Oxo-M: *n* = 6, *N* = 5). ***I***, Steady-state instantaneous firing frequency calculated from responses to a 2x rheobase current injection (control: *n* = 10, *N* =5; Oxo-M: *n* = 3, *N* = 2). ***J***, Maximum instantaneous firing frequency reached during responses to a 2x rheobase current injection (sample sizes are the same as in ***H***). *Statistical analysis,* (***A***) mixed-effects analysis, (***C***-***E***, ***H***) unpaired t-test and (***I-J***) Mann-Whitney tests. * *P* < 0.05, **** *P* < 0.0001; otherwise, not statistically significant.

By 4 dpf, pMNs are able to repetitively fire during sustained depolarization (Gaudreau & Bui, 2024). We therefore investigated whether enhancing *I*_M_ in pMNs with Oxo-M reduced repetitive firing in pMNs at 4 dpf, To do this, we injected current equal to 2x rheobase into 4 dpf pMNs exposed to 20 µM Oxo-M compared their features of repetitive firing to control 4 dpf pMNs (**Fig. 2*F***-***G***). The number of action potentials produced significantly decreased in the presence of Oxo-M relative to control pMNs (control: 32.46 ± 23.85 spikes; Oxo-M: 7.33 ± 3.50 spikes; **Fig. 2*H***) while both steady-state firing frequency (control: 70.73 ± 36.11 Hz; Oxo-M: 95.45 ± 49.66 Hz; **Fig. 2*I***) and maximum instantaneous firing frequency remained unaffected (control: 360.00 ± 95.19 Hz; Oxo-M: 336.00 ±117.50 Hz; **Fig. 2*J***). Overall, these data suggest that the enhancing influence of M2/M4 signaling on *I*_M_ amplitude decreases the ability of pMNs to repetitively fire.

### Dopamine does not alter the properties of I_M_ in pMNs of developing zebrafish

With the finding that muscarine does not inhibit *I*_M_, unlike what has been shown in other species, we sought to determine whether the action of another neurotransmitter had inhibitory influences on *I*_M_. Dopamine (DA) has been identified as a neuromodulator with important influence on the function of spinal locomotor circuits in zebrafish during development (Lambert *et al*., 2012). We aimed to determine whether some of these previously reported effects of DA could be occurring via modulation of *I*_M_ by DA. To this end, we examined possible effects to the properties of *I*_M_ in pMNs exposed to either 10 µM or 100 µM DA at 3, 4, and 5 dpf. At 3 dpf, pMNs exposed to either 10 or 100 µM DA exhibited no changes to the current-voltage relationship of *I*_M_ (**Fig. 3*A-B***), the peak amplitude of *I*_M_ (control: 42.88 ± 18.44 pA; 10 µM dopamine: 44.97 ± 22.80 pA; 100 µM dopamine: 29.94 ± 10.44 pA; **Fig. 3*C***), its activation voltage (control: −54.55 ± 5.68 mV; 10 µM DA: −57.00 ± 4.22 mV; 100 µM DA: −55.45 ± 7.57 mV; **Fig. 3*D***), nor its *V*_1/2_ (control: −47.64 ± 4.31 mV; 10 µM DA: −47.96 ± 6.44 mV; 100 µM DA: −46.04 ± 8.04 mV; **Fig. 3*E***) relative to controls. At 4 dpf, no significant effects by either 10 µM DA or 100 µM DA to the current-voltage relationship of *I*_M_ (**Fig. 3*F-G***), the peak amplitude of *I*_M_ (control: 24.08 ± 10.91 pA; 10 µM DA: 34.25 ± 16.81 pA; 100 µM DA: 22.96 ± 8.82 pA; **Fig. 3*H***), the activation voltage of *I*_M_ (control: −45.63 ± 10.63 mV; 10 µM DA: −53.33 ± 9.35 mV; 100 µM DA: −51.36 ± 6.74 mV; **Fig. 3*I***), or its *V*_1/2_ (control: −41.90 ± 6.75 mV; 10 µM DA: −46.41 ± 7.62 mV; 100 µM DA: −44.44 ± 9.61 mV; **Fig. 3*J***) compared to control pMNs. Similarly, by 5 dpf, neither 10 µM DA nor 100 µM DA altered the current-voltage relationship of *I*_M_ (**Fig. 3*K-L***), the peak amplitude of *I*_M_ (control: 28.63 ± 11.56 pA; 10 µM dopamine: 39.24 ± 9.96 pA; 100 µM dopamine: 23.50 ± 9.62 pA; **Fig. 2*M***), the activation voltage of *I*_M_ (control: −49.09 ± 7.01 mV; 10 µM dopamine: −53.89 ± 5.47 mV; 100 µM dopamine: −50.00 ± 7.07 mV; **Fig. 3*N***) nor its *V*_1/2_ (control: −42.88 ± 9.03 mV; 10 µM dopamine: −44.73 ± 8.20 mV; 100 µM dopamine: −47.79 ± 3.37 mV; **Fig. 3*O***). Overall, these data reveal no effects of dopamine, at both concentrations tested, to the properties of *I*_M_ in 3 to 5 dpf pMNs.

**Figure 3.**
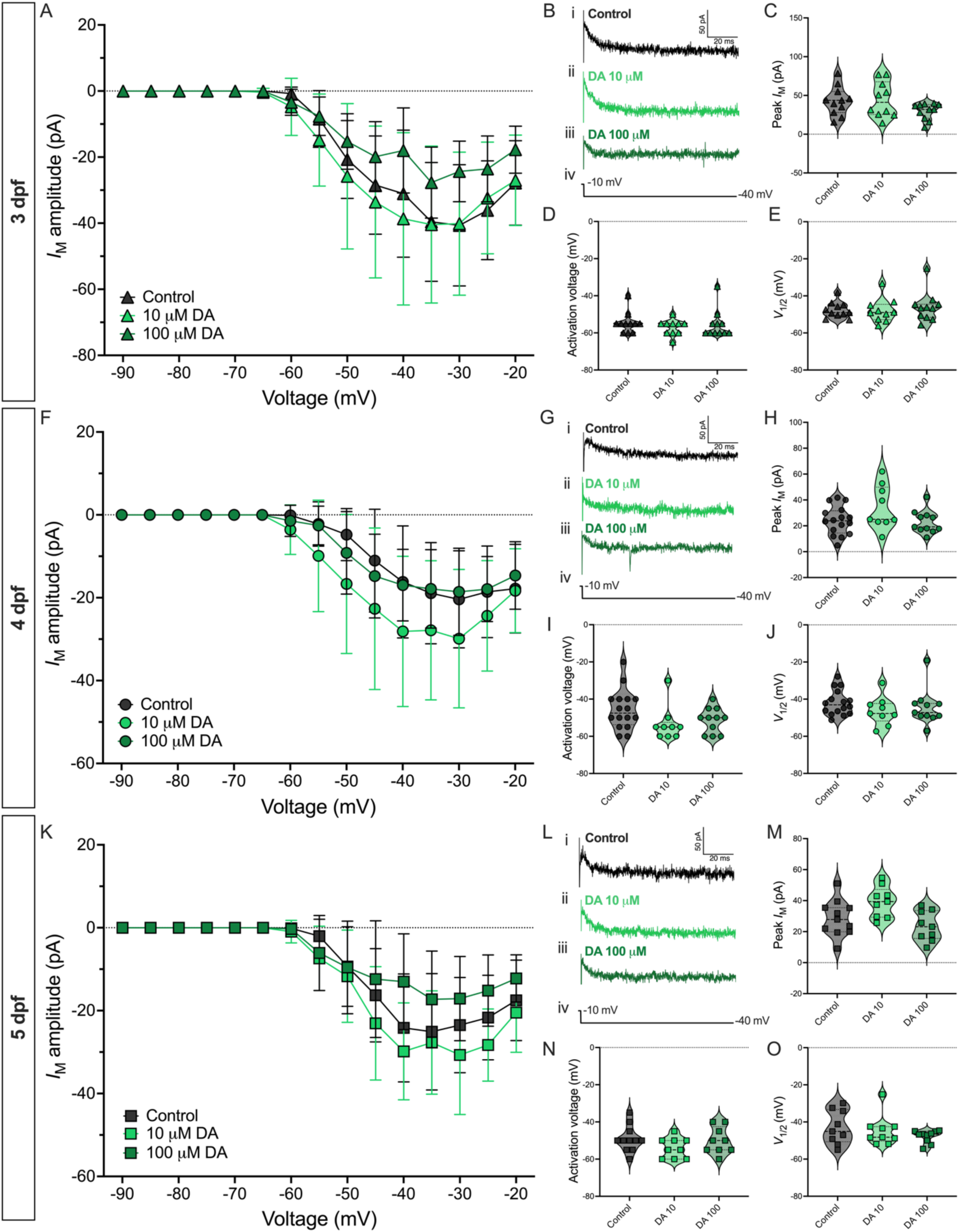
Dopamine does not modulate *I*_M_ in pMNs of developing zebrafish. ***A***, Current-voltage relationship of *I*_M_ in three groups of 3 dpf pMNs: control (*n* = 11, *N =* 6), 10 µM dopamine (DA) (*n* = 10, *N* = 3), 100 µM dopamine (*n* = 11, *N* = 3). ***B***, Examples of current responses from (*i*) control, (*ii*) 10 µM dopamine, and (*iii*) 10 µM dopamine pMNs in response to (*iv*) a −30 mV hyperpolarizing voltage step at 3 dpf. ***C***, Peak amplitude of *I*_M_ across treatment groups at 3 dpf. ***D***, Activation voltage of *I*_M_ across treatment groups at 3 dpf. ***E***, Voltage at which *I*_M_ reaches half its maximal amplitude (*V*_1/2_) across treatment group at 3 dpf. ***F***, Current-voltage relationship of *I*_M_ in three groups of 4 dpf pMNs: control (*n* = 16, *N =* 6), 10 µM dopamine (*n* = 9, *N* = 3), 100 µM dopamine (*n* = 11, *N* = 3). ***G***, Examples of current responses from (*i*) control, (*ii*) 10 µM dopamine, and (*iii*) 100 µM dopamine pMNs in response to (*iv*) a −30 mV hyperpolarizing voltage step at 4 dpf. ***H***, Peak amplitude of *I*_M_ across the treatment groups at 4 dpf. ***I***, Activation voltage of *I*_M_ across treatment groups at 4 dpf. ***J***, Voltage at which *I*_M_ reaches half its maximal amplitude (*V*_1/2_) across treatment group at 4 dpf. ***K***, Current-voltage relationship of *I*_M_ in three groups of 5 dpf pMNs: control (*n* = 11, *N =* 5), 10 µM dopamine (*n* = 9, *N* = 5), 100 µM dopamine (*n* = 9, *N* = 3). ***L***, Examples of current responses from (*i*) control, (*ii*) 10 µM dopamine, and (*iii*) 100 µM dopamine pMNs in response to (*iv*) a −30 mV hyperpolarizing voltage step at 5 dpf. ***M***, Peak amplitude of *I*_M_ across the treatment groups at 5 dpf. ***N***, Activation voltage of *I*_M_ across treatment groups at 5 dpf. ***O***, Voltage at which *I*_M_ reaches half its maximal amplitude (*V*_1/2_) across treatment group at 5 dpf. *Statistical analysis,* (***A***, ***F***, ***K***) mixed-effects analysis, (***D***-***E***, ***I***-***J***, ***M***, ***O***) Kruskal-Wallis test with Dunn’s test for multiple comparisons, and (***C***, ***H****, **N***) ordinary one-way ANOVA with Dunnett’s test for multiple comparisons.

### Serotonin enhances I_M_ in pMNs at 4 dpf

Serotonin (5-HT) has been shown to inhibit *I*_M_ in pyramidal neurons of the weakly electric fish (Deemyad *et al*., 2011) as well as neurons of the hypothalamus in mice (Roepke *et al*., 2012). We therefore decided to test whether exogenous application of 5-HT at 20 µM and 100 µM altered properties of *I*_M_ in pMNs of larval zebrafish (**Fig. 4**). At 3 dpf, no differences between the current-voltage relationship of *I*_M_ (**Fig. 4*A-B***), the peak amplitude of *I*_M_ (control: 42.88 ± 18.44 pA; 20 µM 5-HT: 47.95 ± 22.16 pA; 100 µM 5-HT: 35.07 ± 13.97 pA; **Fig. 4*C***), the activation voltage of *I*_M_ (control: −54.55 ± 5.68 mV; 20 µM 5-HT: −57.50 ± 5.40 mV; 100 µM 5-HT: −55.00 ± 7.50 mV; **Fig. 4*D***) or its *V*_1/2_ (control: −37.64 ± 4.31 mV; 20 µM 5-HT: −51.76 ± 3.59 mV; 100 µM 5-HT: −47.92 ± 5.12 mV; **Fig. 4*E***) in pMNs across treatment groups were observed. By 4 dpf, the average amplitude of *I*_M_ across voltages is increased in pMNs by 20 µM 5-HT, but not 100 µM 5-HT, compared to controls (**Fig. 4*F-G***). Similarly, 20 µM 5-HT, but not 100 µM 5-HT, increases the peak amplitude of *I*_M_ measured in pMNs compared to controls (control: 24.08 ± 10.91 pA; 20 µM 5-HT: 42.36 ± 15.31 pA; 100 µM 5-HT: 27.94 ± 15.59 pA; **Fig. 4*H***). The activation voltage of *I*_M_ remains unchanged by 20 µM 5-HT and 100 µM 5-HT compared to controls (control: −45.63 ± 10.63 mV; 20 µM 5-HT: −51.67 ± 6.85 mV; 100 µM 5-HT: −45.42 ± 9.16 mV; **Fig. 4*I***). Neither 20 µM or 100 µM 5-HT altered *V*_1/2_ relative to controls (control: −41.90 ± 6.75 mV; 20 µM 5-HT: −46.51 ± 6.07 mV; 100 µM 5-HT: −39.83 ± 8.21 mV; **Fig. 4*J***). By 5 dpf, there are no differences across treatment groups to the current-voltage relationship of *I*_M_ (**Fig. 4*K-L***), the peak amplitude of *I*_M_ (control: 28.63 ± 11.56 pA; 20 µM 5-HT: 34.95 ± 15.17 pA; 100 µM 5-HT: 37.53 ± 22.82 pA; **Fig. 4*M***), the activation voltage of *I*_M_ (control: −49.09 ± 7.01 mV; 20 µM 5-HT: −51.54 ± 8.26 mV; 100 µM 5-HT: −47.50 ± 3.78 mV; nor *I*_M_’s *V*_1/2_ (control: −47.79 ± 3.37 mV; 20 µM 5-HT: −45.35 ± 7.02 mV; 100 µM 5-HT: −43.06 ± 3.25 mV; **Fig. 4*O***). These findings reveal that 5-HT at 20 µM may have an overall enhancing effect on *I*_M_ that is most prominent at 4 dpf.

**Figure 4.**
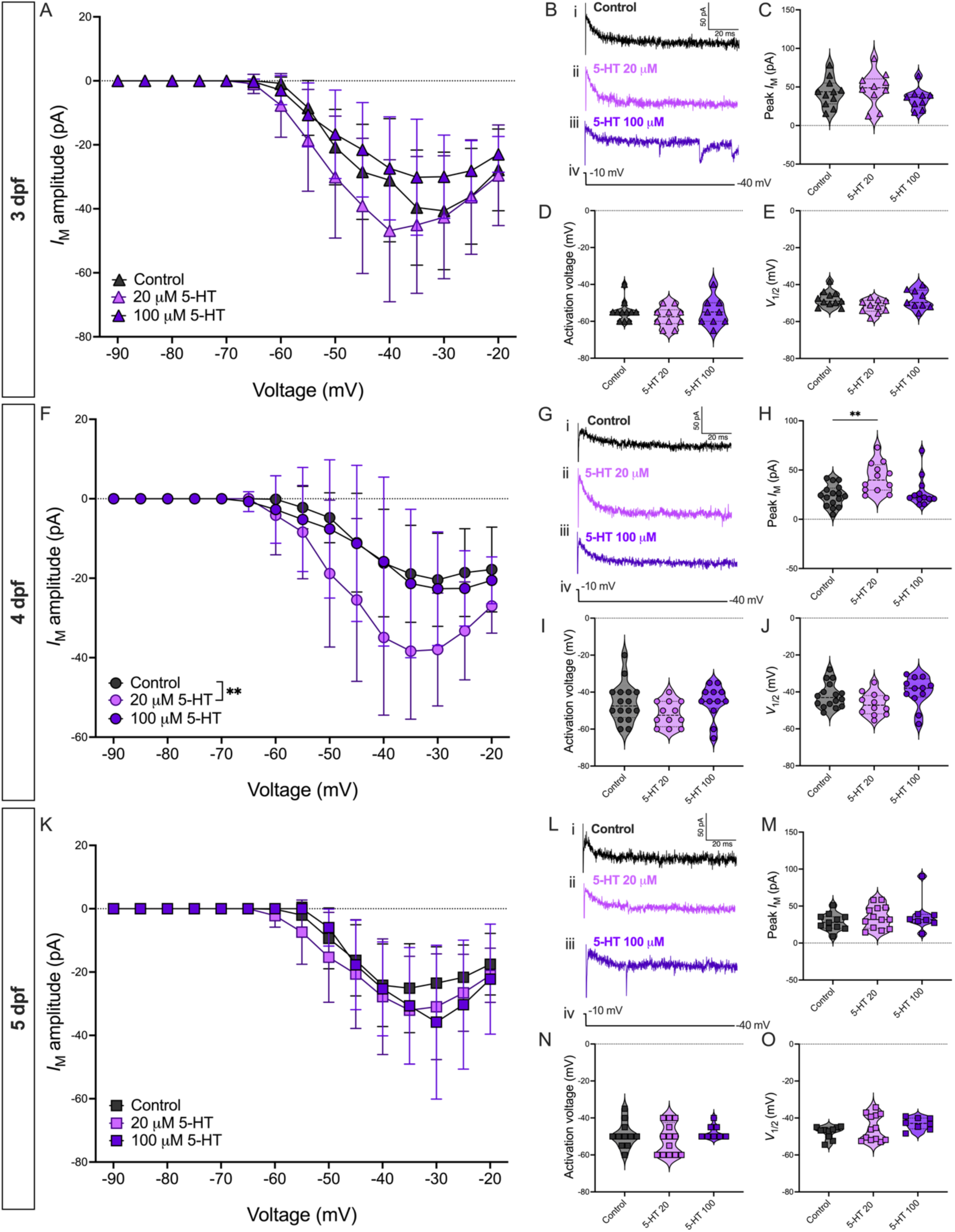
Serotonin enhances *I*_M_ in pMNs of developing zebrafish at 4 dpf. ***A***, Current-voltage relationship of *I*_M_ in three groups of 3 dpf pMNs: control (*n* = 11, *N =* 6), 20 µM serotonin (5-HT) (*n* = 10, *N* = 5), 100 µM 5-HT (*n* = 9, *N* = 5). ***B***, Examples of current responses from (*i*) control, (*ii*) 20 µM 5-HT, and (*iii*) 100 µM 5-HT pMNs in response to (*iv*) a −30 mV hyperpolarizing voltage step at 3 dpf. ***C***, Peak amplitude of *I*_M_ across treatment groups at 3 dpf. ***D***, Activation voltage of *I*_M_ across treatment groups at 3 dpf. ***E***, Voltage at which *I*_M_ reaches half its maximal amplitude (*V*_1/2_) across treatment group at 3 dpf. ***F***, Current-voltage relationship of *I*_M_ in three groups of 4 dpf pMNs: control (*n* = 16, *N =* 6), 20 µM 5-HT (*n* = 12, *N* = 4), 100 µM 5-HT (*n* = 12, *N* = 6). ***G***, Examples of current responses from (*i*) control, (*ii*) 20 µM 5-HT and (*iii*) 100 µM 5-HT pMNs in response to (*iv*) a −30 mV hyperpolarizing voltage step at 4 dpf. ***H***, Peak amplitude of *I*_M_ across the treatment groups at 4 dpf. ***I***, Activation voltage of *I*_M_ across treatment groups at 4 dpf. ***J***, Voltage at which *I*_M_ reaches half its maximal amplitude (*V*_1/2_) across treatment group at 4 dpf. ***K***, Current-voltage relationship of *I*_M_ in three groups of 5 dpf pMNs: control (*n* = 11, *N =* 5), 20 µM 5-HT (*n* = 13, *N* = 5), 100 µM 5-HT (*n* = 8, *N* = 4). ***L***, Examples of current responses from (*i*) control, (*ii*) 20 µM 5-HT, and (*iii*) 100 µM 5-HT pMNs in response to (*iv*) a −30 mV hyperpolarizing voltage step at 5 dpf. ***M***, Peak amplitude of *I*_M_ across the treatment groups at 5 dpf. ***N***, Activation voltage of *I*_M_ across treatment groups at 5 dpf. ***O***, Voltage at which *I*_M_ reaches half its maximal amplitude (*V*_1/2_) across treatment group at 5 dpf. *Statistical analysis,* (***A***, ***F***, ***K***) mixed-effects analysis, (***H***, ***M***-***O***) Kruskal-Wallis test with Dunn’s test for multiple comparisons, and (***D***-***E***, ***I***-***J***) ordinary one-way ANOVA with Dunnett’s test for multiple comparisons: ** *P* < 0.01; otherwise, not statistically significant.

### Serotonin does not modulate I_M_ in pMNs via 5HT_2A_ receptors

We next furthered our investigation into the role of serotonin in the modulation of *I*_M_ in pMNs of developing zebrafish by investigating how the selective 5HT_2A_ receptor agonist TCB-2 influences properties of *I*_M_. As with the experiments described above, we compared the properties of *I*_M_ between two distinct groups of pMNs, one being control pMNs and the other being pMNs exposed to 10 µM TCB-2 before recordings began. At 3 dpf, TCB-2 does not alter the current-voltage relationship of *I*_M_ (**Fig. 5*A-B***), its peak amplitude (control: 42.88 ± 18.44 pA; TCB-2: 40.86 ± 17.38 pA; **Fig. 5*C***), activation voltage (control: −54.55 ± 5.68 mV; TCB-2: −55.55 ± 7.70 mV; **Fig. 5*D***), or its *V*_1/2_ (control: −49.39 ± 6.19 mV; TCB-2: −47.64 ± 4.31 mV; **Fig. 5*E***). This is also the case at 4 dpf, where TCB-2 does not significantly influence the current-voltage relationship of *I*_M_ (**Fig. 5*F-G***), its peak amplitude (control: 24.08 ± 10.91 pA; TCB-2: 25.15 ± 9.31 pA; **Fig. 5*H***), activation voltage (control: −45.63 ± 10.63 mV; TCB-2: −45.45 ± 7.23 mV; **Fig. 5*I***), or *V*_1/2_ (control: −41.90 ± 6.75 mV; TCB-2: −43.34 ± 6.04 mV; **Fig. 5*J***). Similarly, by 5 dpf, the current-voltage relationship (**Fig. 5*K-L***), peak amplitude (control: 28.63 ± 11.56 pA; TCB-2: 29.26 ± 9.67 pA; **Fig. 5*M***), activation voltage (control: −49.09 ± 7.01 mV; TCB-2: −49.09 ± 7.36 mV; **Fig. 5*N***), and *V*_1/2_ (control: −47.79 ± 3.37 mV; TCB-2: −44.45 ± 7.06 mV; **Fig. 5*O***) of *I*_M_ are not altered by TCB-2 relative to controls. These results demonstrate that serotonin via 5HT_2A_ receptors does not modulate *I*_M_ in pMNs from 3 to 5 dpf.

**Figure 5.**
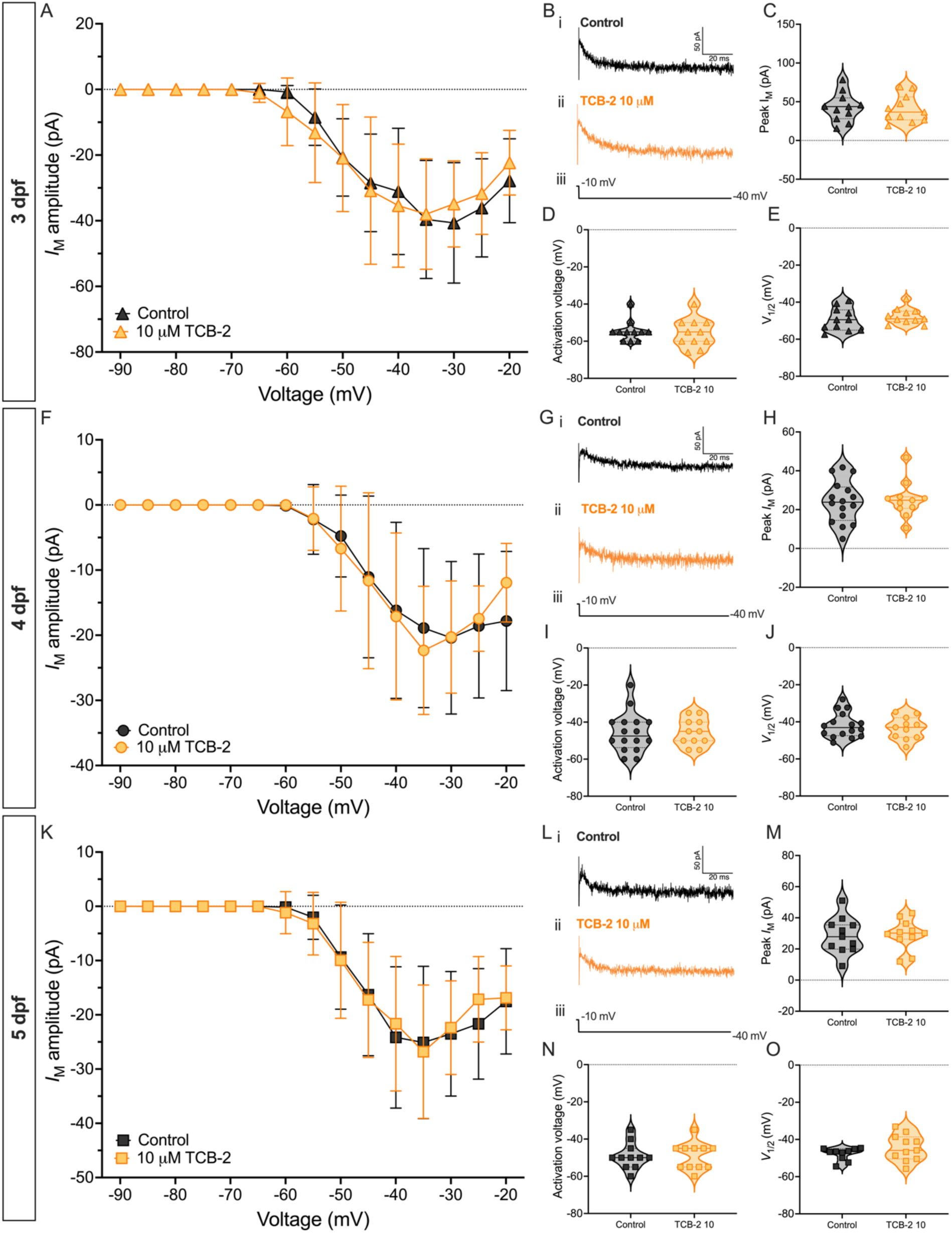
5HT_2A_ agonist does not modulate *I*_M_ in pMNs during zebrafish development. ***A***, Current-voltage relationship of *I*_M_ in two groups of 3 dpf pMNs: control (*n* = 11, *N =* 6) and 10 µM TCB-2 (*n* = 11, *N* = 5). ***B***, Examples of current responses from (*i*) control and (*ii*) 10 µM TCB-2 pMNs in response to (*iii*) a −30 mV hyperpolarizing voltage step at 3 dpf. ***C***, Peak amplitude of *I*_M_ across treatment groups at 3 dpf. ***D***, Activation voltage of *I*_M_ across treatment groups at 3 dpf. ***E***, Voltage at which *I*_M_ reaches half its maximal amplitude (*V*_1/2_) across treatment group at 3 dpf. ***F***, Current-voltage relationship of *I*_M_ in two groups of 4 dpf pMNs: control (*n* = 16, *N =* 6), and 10 µM TCB-2 (*n* = 11, *N* = 4). ***G***, Examples of current responses from (*i*) control and (*ii*) 10 µM TCB-2 pMNs in response to (*iii*) a −30 mV hyperpolarizing voltage step at 4 dpf. ***H***, Peak amplitude of *I*_M_ across the treatment groups at 4 dpf. ***I***, Activation voltage of *I*_M_ across treatment groups at 4 dpf. ***J***, Voltage at which *I*_M_ reaches half its maximal amplitude (*V*_1/2_) across treatment group at 4 dpf. ***K***, Current-voltage relationship of *I*_M_ in two groups of 5 dpf pMNs: control (*n* = 11, *N =* 5) and 10 µM TCB-2 (*n* = 11, *N* = 5). ***L***, Examples of current responses from (*i*) control and (*ii*) 10 µM TCB-2 pMNs in response to (*iii*) a −30 mV hyperpolarizing voltage step at 5 dpf. ***M***, Peak amplitude of *I*_M_ across the treatment groups at 5 dpf. ***N***, Activation voltage of *I*_M_ across treatment groups at 5 dpf. ***O***, Voltage at which *I*_M_ reaches half its maximal amplitude (*V*_1/2_) across treatment group at 5 dpf. *Statistical analysis,* (***A***, ***F***, ***K***) mixed-effects analysis, (***C***, ***H***-***J***, ***M***-***N***) unpaired t-tests, and (***D***, ***O***) Mann-Whitney tests.

### Serotonin via 5HT_1A_ receptors inhibits I_M_ in pMNs at 3 dpf

Since our TCB-2 experiments revealed no modulation of *I*_M_ in pMNs by serotonin via 5HT_2A_ receptors, we next investigated serotonin’s possible influence on *I*_M_ via 5HT_1A_ receptors. We compared the properties of *I*_M_ in pMNs exposed to 10 µM 8-OH-DPAT to those of control pMNs at 3, 4, and 5 dpf. At 3 dpf, we find that 8-OH-DPAT inhibits *I*_M_ as is observed by a significant decrease in the average amplitude of *I*_M_ across voltages (**Fig. 6*A-B***) and the peak amplitude of *I*_M_ (control: 42.88 ± 18.44 pA; 10 µM 8-OH-DPAT: 22.94 ± 11.05 pA; **Fig. 6*C***). *I*_M_’s activation voltage (control: −54.44 ± 5.68 mV; 10 µM 8-OH-DPAT: −52.50 ± 10.34 mV; **Fig. 6*D***) and *V*_1/2_ (control: −47.64 ± 4.31 mV; 10 µM 8-OH-DPAT: −42.29 ± 7.37 mV; **Fig. 6*E***) remained unchanged by 8-OH-DPAT in pMNs at 3 dpf. At 4 dpf, 8-OH-DPAT does not alter the current-voltage relationship of *I*_M_ (**Fig. 6*F-G***), the peak amplitude of *I*_M_ (control: 24.08 ± 10.91 pA; 10 µM 8-OH-DPAT: 31.62 ± 26.83 pA; **Fig. 6*H***), the activation voltage of *I*_M_ (control: −45.63 ± 10.63 mV; 10 µM 8-OH-DPAT: −50.91 ± 9.44 mV; **Fig. 6*I***), nor *I*_M_’s *V*_1/2_ (control: −41.90 ± 6.75 mV; 10 µM 8-OH-DPAT: −45.42 ± 10.78 mV; **Fig. 6*J***) in pMNs compared to controls. Similarly, at 5 dpf, 8-OH-DPAT does not alter the current-voltage relationship of *I*_M_ (**Fig. 6*K-L***), the peak amplitude of *I*_M_ (control: 28.63 ± 11.56 pA; 10 µM 8-OH-DPAT: 34.14 ± 15.99 pA; **Fig. 6*M***), nor *I*_M_’s *V*_1/2_ (control: −47.79 ± 3.37 mV; 10 µM 8-OH-DPAT: −49.87 ± 3.66 mV; **Fig. 6*O***) in pMNs compared to controls. TCB-2 does however hyperpolarize the activation voltage of *I*_M_ (control: −49.09 ± 7.01 mV; 10 µM 8-OH-DPAT: −55.56 ± 4.64 mV; **Fig. 6*N***). These data support an inhibitory influence of serotonin signaling via 5-HT_1A_ receptors on *I*_M_ specifically at 3 dpf.

**Figure 6.**
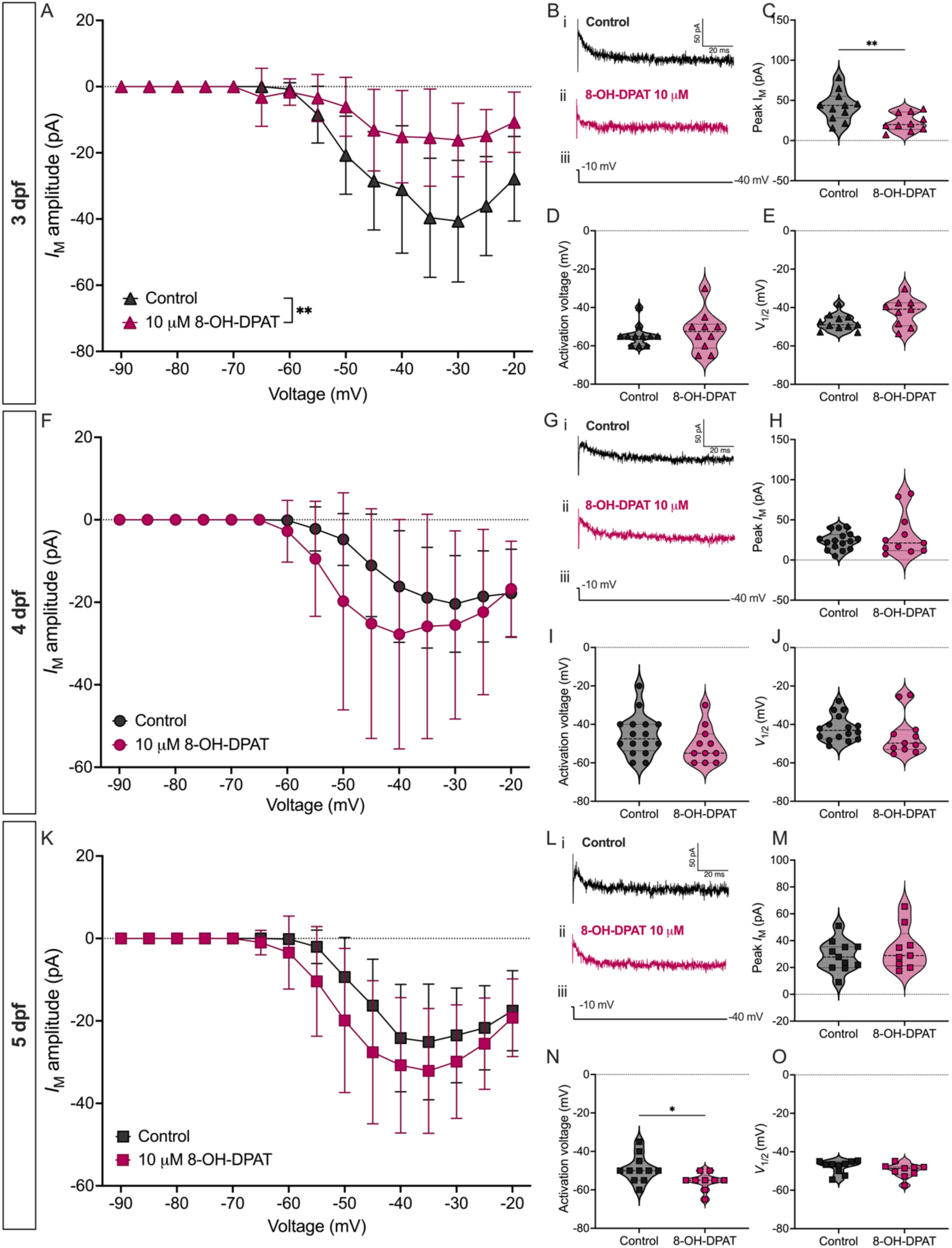
5HT_1A_ agonist inhibits *I*_M_ in pMNs at 3 dpf. ***A***, Current-voltage relationship of *I*_M_ in two groups of 3 dpf pMNs: control (*n* = 11, *N =* 6) and 10 µM 8-OH-DPAT (*n* = 10, *N* = 4). ***B***, Examples of current responses from (*i*) control and (*ii*) 10 µM 8-OH-DPAT pMNs in response to (*iii*) a −30 mV hyperpolarizing voltage step at 3 dpf. ***C***, Peak amplitude of *I*_M_ across treatment groups at 3 dpf. ***D***, Activation voltage of *I*_M_ across treatment groups at 3 dpf. ***E***, Voltage at which *I*_M_ reaches half its maximal amplitude (*V*_1/2_) across treatment group at 3 dpf. ***F***, Current-voltage relationship of *I*_M_ in two groups of 4 dpf pMNs: control (*n* = 16, *N =* 6), and 10 µM 8-OH-DPAT (*n* = 11, *N* = 5). ***G***, Examples of current responses from (*i*) control and (*ii*) 10 µM 8-OH-DPAT pMNs in response to (*iii*) a −30 mV hyperpolarizing voltage step at 4 dpf. ***H***, Peak amplitude of *I*_M_ across the treatment groups at 4 dpf. ***I***, Activation voltage of *I*_M_ across treatment groups at 4 dpf. ***J***, Voltage at which *I*_M_ reaches half its maximal amplitude (*V*_1/2_) across treatment group at 4 dpf. ***K***, Current-voltage relationship of *I*_M_ in two groups of 5 dpf pMNs: control (*n* = 11, *N =* 5) and 10 µM 8-OH-DPAT (*n* = 9, *N* = 4). ***L***, Examples of current responses from (*i*) control and (*ii*) 10 µM 8-OH-DPAT pMNs in response to (*iii*) a −30 mV hyperpolarizing voltage step at 5 dpf. ***M***, Peak amplitude of *I*_M_ across the treatment groups at 5 dpf. ***N***, Activation voltage of *I*_M_ across treatment groups at 5 dpf. ***O***, Voltage at which *I*_M_ reaches half its maximal amplitude (*V*_1/2_) across treatment group at 5 dpf. *Statistical analysis,* (***A***, ***F***, ***K***) mixed-effects analysis, (***C***, ***E***, ***I***, ***M***-***N***) unpaired t-tests, and (***D***, ***H***, ***J***, ***O***) Mann-Whitney tests: * *P* < 0.05, ** *P* < 0.01; otherwise, not statistically significant.

### Serotonin signaling via 5HT_1A_ receptors increases repetitive firing in pMNs at 3 dpf

We have previously shown that inhibiting *I*_M_ with 10 µM XE-991 in pMNs increased their ability to repetitively fire at 3 dpf, the age at which *I*_M_ appears to be largest in magnitude during early development (Gaudreau & Bui, 2024). Having identified 5HT_1A_ receptors as mediating serotonin’s inhibitory influence over *I*_M_, we aimed to investigate whether 8-OH-DPAT increased the ability of 3 dpf pMNs to repetitively fire in a manner similar to application of the *I*_M_-specific inhibitor XE-991. To do this, we injected 3 dpf pMNs exposed to 10 µM 8-OH-DPAT with current equal to 2x rheobase and compared their features of repetitive firing to control 3 dpf pMNs (**Fig. 7*A***-***B***). We find that 8-OH-DPAT significantly increases the number of spikes generated (control: 8.33 ± 6.85 spikes; 8-OH-DPAT: 23.00 ± 13.38 spikes; **Fig. 7*C***) as well as the steady-state firing frequency (control: 29.73 ± 3.71 Hz; 8-OH-DPAT: 58.36 ± 22.83 Hz; **Fig. 7*D***) while the maximum instantaneous firing frequency remained unchanged (control: 392.2 ± 185.0 Hz; 8-OH-DPAT: 337.3 ± 94.2 Hz; **Fig. 7*E***) relative to control pMNs. Overall, these results suggest that serotonin via 5HT_1A_ receptors can promote repetitive firing in pMNs via inhibition of *I*_M_ at 3 dpf.

**Figure 7.**
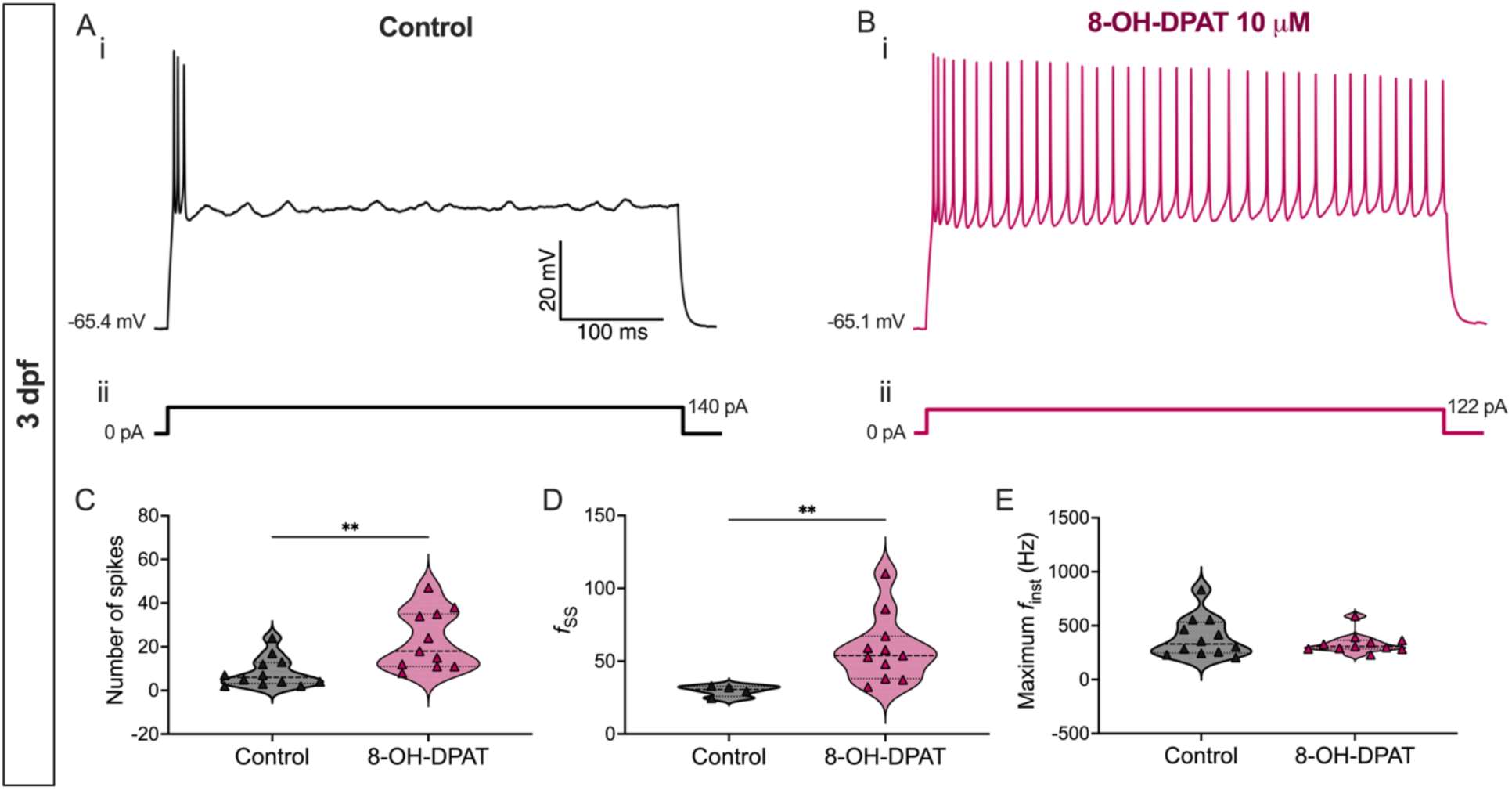
5-HT_1A_ agonist increases repetitive firing in pMNs at 3 dpf. ***A***-***B***, Representative current-clamp responses of (***Ai***) control pMNs and (***Bi***) those exposed to 10 μM 8-OH-DPAT in response to (***ii***) a current injection equal to 2x rheobase. ***C***, Number of spikes generated in response to a 2x rheobase current injection (control: *n* = 12, *N* = 6; 8-OH-DPAT: *n* = 11, *N* = 5). ***D***, Steady-state instantaneous firing frequency calculated from responses to a 2x rheobase current injection (sample sizes are the same as in ***C***). ***E***, Maximum instantaneous firing frequency reached during responses to a 2x rheobase current injection (control: *n* = 4, *N* = 2; 8-OH-DPAT: *n* = 11, *N* = 5). *Statistical analysis,* Mann-Whitney tests. ** *P* < 0.01; otherwise, not statistically significant.

## DISCUSSION

Neuromodulators such as DA and 5-HT are known to influence motor maturation, as evidenced by work in the developing zebrafish (Brustein *et al*., 2003; Lambert *et al*., 2012). Specific neuronal targets of this modulation however remain largely unknown as do the mechanism by which neuromodulators exert their influence on neurons. In this study, we investigated the subthreshold non-inactivating potassium current, *I*_M_, as a possible target for neuromodulation in spinal locomotor circuits of the developing zebrafish. *I*_M_ shows non-monotonic developmental dynamics in pMNs during a time period where zebrafish transition from crude, ballistic coiling movements to more refined swimming maneuvers characterized by low-amplitude repetitive tail beats (Gaudreau & Bui, 2024). At the same time, the firing properties of pMNs mature with decreasing spike frequency adaptation and increased sustained firing (Gaudreau & Bui, 2024). We reasoned that neuromodulation of *I*_M_ may also shift during this time period of rapid changes in the amplitude of this current. We characterized the influence of muscarine, DA, and 5-HT on the activity of *I*_M_ in primary motoneurons from 3 to 5 dpf as developing zebrafish undergo motor maturation (summarized in **Fig. 8**).

**Figure 8.**
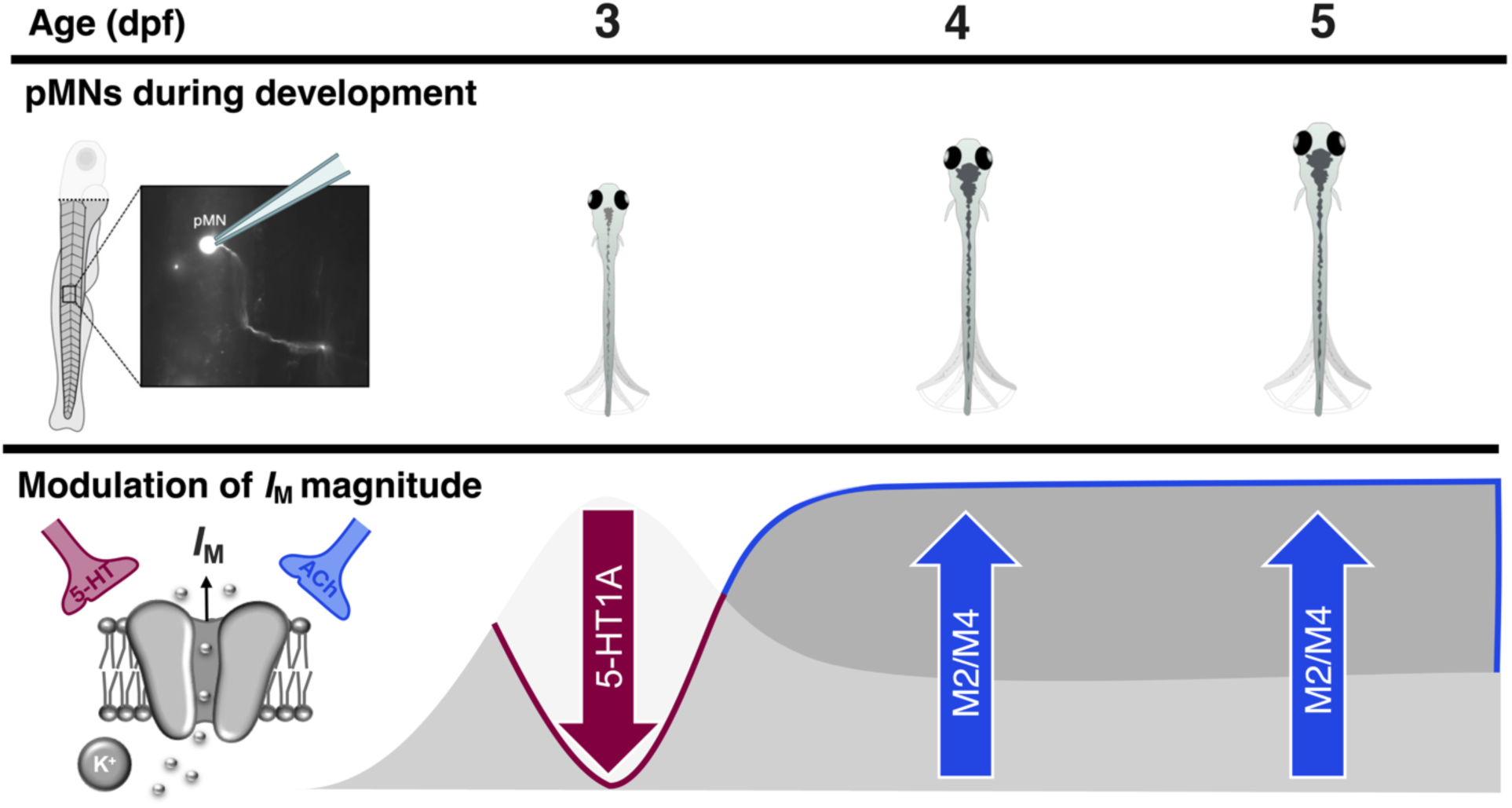
Distinct modulatory influence of serotonin and acetylcholine on *I*_M_ in pMNs during zebrafish development. At 3 dpf, when the magnitude of *I*_M_ peaks (Gaudreau & Bui, 2024) during early zebrafish development, serotonin signaling via 5HT_1A_ receptors acts to inhibit *I*_M_. By 3 dpf, when the magnitude of *I*_M_ has subsequently decreased, equivalent activation of 5HT_1A_ receptors no longer inhibits *I*_M_. Instead, we find that at this age, *I*_M_ is subject to enhancement by acetylcholine via M2/M4 receptors. *I*_M_ in pMNs may present itself as a dial for neuromodulation to fine-tune pMN excitability and firing activity during zebrafish development.

### Muscarine as an I_M_ enhancer in pMNs of developing zebrafish

*I*_M_ was designated as the M-current based on the finding that it was inhibited by muscarine at the time of its discovery (Brown & Adams, 1980). Because of this, we had expected to see inhibition of *I*_M_ in pMNs exposed to muscarine. Intriguingly however, we found that muscarine instead had an enhancing effect on *I*_M_ in pMNs at 4 and 5 dpf with no effects observed at 3 dpf. Traditional inhibition of *I*_M_ by muscarine is mediated by muscarinic acetylcholine receptors (mAChRs) coupled to G_q/11_ proteins whose activation leads to the eventual depletion of phosphatidylinositol 4,5-biphosphate (PIP_2_) at the membrane which directly influences the activity of Kv7.2/7.3 channels that mediate *I*_M_ (Zhang *et al*., 2003; Suh & Hille, 2007). A study in neurons of the mouse dentate gyrus however reveals that, contrary to most other findings, acetylcholine signaling via M_1_ receptors actually enhances the activity of *I*_M_ by increasing levels of PIP_2_ (Carver & Shapiro, 2019). In line with these findings, our data support the idea that cholinergic influence over levels of PIP_2_ through muscarinic receptors, and subsequently *I*_M_ activity is neuron-specific and can deviate from the usual suppression of this current. We demonstrate that the preferential M2/M4 mAChR agonist oxotremorine-M enhances *I*_M_ in 4 dpf pMNs, raising the possibility that in these neurons M2 or M4 mAChR activation may ultimately lead to increased PIP_2_ levels. Alternatively, this enhancement of *I*_M_ could be brought on by a PIP_2_-independent mechanism (Marrion, 1997). Regardless of the precise mAChRs and mechanisms involved, our data reveals functional impact of *I*_M_ enhancement by M2/M4 activation wherein oxotremorine-M limits the ability of pMNs to repetitively fire. Activation of M2 mAChRs in fast motoneurons of the adult zebrafish also limits their spiking activity (Bertuzzi & Ampatzis, 2018). Our data raises the possibility that *I*_M_ in fast motoneurons could continue to be the target of cholinergic influence on decreased repetitive firing in the adult zebrafish. Further studies are necessary to uncover both the source of acetylcholine – cholinergic spinal V2a (Guan *et al*., 2021) and V0c neurons being prime candidates (Kelly *et al*., 2023) – and the contexts in which mAChRs might be activated to enhance *I*_M_, and consequently limit firing, in pMNs.

### Serotonin as both an enhancer and inhibitor of I_M_ in pMNs of developing zebrafish

Exogenous application of 5-HT leads to obvious enhancing of *I*_M_ in pMNs at 4 dpf, but not at 3 or 5 dpf. When we specifically target 5-HT_1A_ receptors using the selective agonist 8-OH-DPAT, however, we find that *I*_M_ is inhibited in pMNs at 3 dpf. Several instances of opposing roles of 5-HT on the excitability of individual neurons have been observed in other systems. For example, 5-HT_2_ receptor mediated serotonergic signaling caused membrane depolarization while 5-HT_1A_ receptor mediated serotonergic signaling resulted in hyperpolarization in pyramidal cells of the cortex in rats (Araneda & Andrade, 1991). In turtle motoneurons, application of 5-HT to the dendritic tree promoted plateau potentials through 5-HT_2A_ activation whereas application to the perisomatic region sometimes inhibited spike generation through 5-HT_1A/7_ receptors (Perrier & Hounsgaard, 2003; Perrier & Cotel, 2008). This raises the intriguing possibility that the lack of effect of 5-HT on *I*_M_ in 3 dpf pMN despite the inhibition observed by activation of 5-HT_1A_ receptors through 8-OH-DPAT administration is due to the presence of a different 5-HT receptor subtype that increases *I*_M_ at that age, and that these two receptors are distributed at different locations. Electrophysiology experiments testing the effects of additional 5-HT receptor subtypes, other than 5HT_1A_ and 5HT_2A_, on the properties of *I*_M_ in pMNs would reveal which receptor is responsible for the 5-HT-driven enhancement of *I*_M_ that we have observed, further delineating the mechanisms underlying *I*_M_ modulation.

### Concentration-dependent neuromodulation of spinal locomotor networks

The presence of multiple receptor subtypes on a single neuron can help explain concentration-dependent effects that we observed. In our investigation into the effects of muscarine, DA, and 5-HT, on the modulation of *I*_M_, we bath-applied each neuromodulator at relatively low and high concentrations. 5-HT and muscarine both changed the magnitude of *I*_M_ at the lower concentrations – 20 and 10 μM, respectively – while showing no effects at 100 μM relative to controls. With distinct receptors having varying affinities for their respective neuromodulators, the ultimate effect to *I*_M_ in pMNs is likely to depend both on the identity of receptors expressed and the concentration of neuromodulator present. Indeed, modulatory action of DA on the activity of rat hypothalamus neurons is concentration-dependent, with low versus high levels of DA determining the direction of dopaminergic modulation (Linehan *et al*., 2015). Opposing effects of DA in the striatum (Stoof & Kebabian, 1981) has long been shown to affect motor control (Kravitz *et al*., 2010). This type of dual complimentary action of neuromodulators has also been observed within spinal locomotor networks across species. ACh, DA, and 5-HT can each exert opposing or concentration-dependent influences on motor output that is dependent on the receptor subtypes activated within each neuromodulator system (Harris-Warrick & Cohen, 1985; Clemens *et al*., 2012; Sharples *et al*., 2015; Nascimento *et al*., 2019). At the motoneuron level, it has been demonstrated in the adult zebrafish that M2 activation decreases firing while activation of other mAChRs increases firing (Bertuzzi & Ampatzis, 2018). We therefore posit that the observed lack of effect of higher doses of serotonin and muscarine can be attributed to the broad activation of receptor subtypes that may ultimately mask effects brought on by activation of higher affinity receptors at lower neuromodulator concentrations. Identification and precise pharmacological targeting of specific neuromodulator receptor subtypes expressed in pMNs is necessary to delineate concentration-dependent effects of neuromodulators to the magnitude of *I*_M_ in these neurons.

### Developmental changes to neuromodulatory influence on I_M_ in pMNs of developing zebrafish

In addition to having uncovered neuromodulators that differentially modulate the activity of *I*_M_, we have revealed age-related differences in the effects of these neuromodulators during the 3 to 5 dpf time window. We show that muscarine enhances *I*_M_ at 4 and 5 dpf, but not at 3 dpf. This is particularly intriguing as this coincides with the reduction in the magnitude of *I*_M_ we have observed from 3 dpf to 4 and 5 dpf. It appears as though when *I*_M_ is already prominent in pMNs at 3 dpf, there is no mAChR-mediated enhancement of *I*_M_. Furthermore, we show that 5-HT_1A_ receptor agonist 8-OH-DPAT inhibits *I*_M_, but only at 3 dpf when the magnitude of *I*_M_ in pMNs is largest during the investigated time period. In this case, it appears that when basal levels of *I*_M_ are low, as they are at 4 and 5 dpf, there is no 5-HT_1A_ mediated inhibition of *I*_M_. These findings support the idea that different neuromodulator systems work together to fine-tune basal activity levels of *I*_M_ that change during development. At 3 dpf when larvae are largely inactive, *I*_M_ in pMNs is large. Inhibition of *I*_M_ by 5-HT_1A_ signaling may help recruit pMNs for fast movements by alleviating the control *I*_M_ has on pMN excitability and repetitive firing. Acetylcholine via mAChRs may only come into play as an enhancer of *I*_M_ by 4-5 dpf because there may be no need for further enhancement of *I*_M_ at 3 dpf. By 4-5 dpf when larvae are more active, *I*_M_ is reduced in magnitude in pMNs compared to 3 dpf. At this point, pMNs may benefit from *I*_M_ enhancement by ACh in contexts where pMN excitability and/or repetitive firing should be limited, for example during transitions from fast to slow swims when pMNs are no longer needed. Whether the push-pull effects that serotonin and acetylcholine exert on *I*_M_ at different ages, which seems to depend on when the amplitude of this current is high or low, relies on the same biochemical pathway (e.g. PIP_2_ signaling) or separate pathways, remains to be determined.

### Behavioural contexts of I_M_ neuromodulation in pMNs

We have shown that *I*_M_ in pMNs is a target of neuromodulation by 5-HT and ACh. Putatively through enhancement and inhibition of *I*_M_, 5-HT and ACh increases and limits, respectively, the ability of pMNs to repetitively fire. This raises the question: in what contexts would *I*_M_ in pMNs need to be modulated? pMNs are recruited primarily during fast and crude movements such as coilings, escapes, fast swims, and struggling (Liu & Westerfield, 1988; Liao & Fetcho, 2008; McLean & Fetcho, 2009; Tong & McDearmid, 2012; Wen *et al*., 2020). Neuromodulation by DA has been shown to prime pMNs for involvement in optomotor responses through modulation of their intrinsic properties (Jha & Thirumalai, 2020). Therefore, neuromodulation of *I*_M_ could also help ensure the degree to which pMNs repetitively fire during various movements, as evidenced by the effect of pharmacological modulation of this current on escapes and swimming (Gaudreau & Bui, 2025). For example, ACh could enhance *I*_M_ to limit pMN firing during the fastest swims requiring high-frequency tail beats while 5-HT could inhibit *I*_M_ to promote repetitive firing in the case of struggling when long-lasting muscle contraction is required (Liao & Fetcho, 2008). Neuromodulation of *I*_M_ in pMNs may supplement circuit connectivity to ensure smooth transitions between swim speeds and appropriate muscle contraction for effective struggling. Indeed, neuromodulation may be more powerful than synaptic inputs in influencing motoneuron excitability and overall motor output (Heckman *et al*., 2009).

To conclude, we have revealed how different neuromodulators influence pMN firing through *I*_M_. We identify two neuromodulator systems that serve as a means to regulate the activity of pMNs in opposite directions via modulation of *I*_M_ and also reveal how their respective influences on *I*_M_ varies within a short developmental time window depending on the age-dependent fluctuations of this current. This work highlights how distinct neuromodulators can exert complimentary and dynamic influences on the activity of ion currents during development, perhaps to facilitate changes to spinal locomotor circuits during motor maturation.

## DATA AVAILABILITY

Data will be made available upon request.

## ACKNOWLEDGMENTS

We would like to thank John Lewis, Michael Jonz, Emily Standen, and Minoru Koyama for their valuable input during the study. This work was supported by the Natural Sciences and Engineering Research Council of Canada (NSERC): NSERC Discovery Grant, Grant Number: RGPIN-2022-03898 (to TVB); NSERC Canadian Graduate Scholarship M award, Award Number: NSERC 553401-2020 (to SFG); NSERC Postgraduate Scholarship D, Award Number: 569969-2022 (to SFG).

## AUTHOR CONTRIBUTIONS

SFG: Conceived and designed research, performed experiments, analyzed data, interpreted results of experiments, prepared figures, drafted manuscript, edited and revised manuscript.

TVB: Conceived and designed research, drafted manuscript, edited and revised manuscript, and approved final version of the manuscript.

